# Serine ubiquitination of p62 regulates Nrf2 dependent redox homeostasis

**DOI:** 10.1101/2023.06.17.545420

**Authors:** Rukmini Mukherjee, Marina Hoffmann, Alexis Gonzalez, Anshu Bhattacharya, Santosh Kumar Kuncha, Rajeshwari Rathore, Donghyuk Shin, Thomas Colby, Ivan Matic, Mohit Misra, Ivan Dikic

## Abstract

The KEAP1–Nrf2 axis is essential for the cellular response against metabolic and oxidative stress. KEAP1 is an adaptor protein of Cullin-3 ubiquitin ligase that controls the cellular levels of Nrf2, a critical transcription factor of several cytoprotective genes. Oxidative stress, defective autophagy and pathogenic infections activate Nrf2 signaling through phosphorylation of the adaptor protein p62, which competes with Nrf2 for binding to KEAP1. Here we show that phosphoribosyl-linked serine ubiquitination of p62 catalyzed by SidE effectors of *Legionella pneumophila* controls Nrf2 signaling and cell metabolism upon *Legionella* infection. Serine ubiquitination of p62 sterically blocks its binding to KEAP1, resulting in Nrf2 ubiquitination and degradation. This reduces Nrf2-dependent antioxidant synthesis in the early phase of infection. Levels of serine ubiquitinated p62 diminish in the later stage of infection allowing the expression of Nrf2-target genes; resulting in a differential regulation of the host metabolome and proteome in a Nrf2 dependent manner.

## Introduction

Reactive oxygen species (ROS) are important to maintain redox homeostasis in the cell. Free radicals like hydroxyl (OH•), superoxide (O2•–), nitric oxide (NO•), nitrogen dioxide (NO2•), peroxyl (ROO•) and lipid peroxyl (LOO•) are generated in enzymatic reactions in cellular compartments which alter the redox state of the cell. Oxidative stress arising from high ROS levels activate redox-sensitive transcriptional factors including the nuclear factor-erythroid 2– related factor 2 (Nrf2) which produce antioxidants to restore the redox state of the cell. Under homeostatic conditions, nuclear levels of Nrf2 are kept low by its cytosolic repressor KEAP1 which is a substrate adapter protein for the Cullin 3 (CUL3)-ring box 1 (RBX1) E3-ubiquitin ligase system. KEAP1 targets Nrf2 for ubiquitination and proteasomal degradation. Under oxidative stress, specific cysteines on KEAP1 are oxidised, leading to conformational changes which release the KEAP1-bound Nrf2, allowing it to translocate to the nucleus where it heterodimerizes with a small musculoaponeurotic fibrosarcoma (sMAF) protein, binds to promoters with antioxidant response elements (ARE) to drive the transcription of Nrf2-target genes (Kobayashi et al., 2004).

Redox signaling is closely associated with cellular metabolism and autophagy. The direct interaction between the autophagy adaptor protein p62/SQSTM1 and KEAP1 links autophagy to Nrf2 signaling. p62/SQSTM1 is a multifunctional protein with several domains, including a Phox1 and Bem1p (PB1) domain, a zinc finger (ZZ), two nuclear localization signals (NLSs), a TRAF6 binding (TB) domain, a nuclear export signal (NES), an LC3-interacting region (LIR), a Keap1-interacting region (KIR), and an ubiquitin-associated (UBA) domain. p62 is well-known for its role in selective autophagy of ubiquitinated cargo comprising intracellular bacteria, depolarized mitochondria or ubiquitinated protein aggregates (Danieli and Martens, 2018, Herhaus and Dikic, 2018, Sanchez-martin et al., 2019). Binding of p62 to ubiquitinated cargo is facilitated by phosphorylation of specific serine residues in its UBA domain (S403, S407). Upon binding to ubiquitinated cargo, S351 within the KIR motif of p62 gets phosphorylated, increasing the binding affinity of its KIR motif to KEAP1. KEAP1 binding to p62 destabilises the KEAP1-Nrf2 interaction, causing Nrf2 stabilisation and its nuclear translocation (Komatsu et al., 2010, Ichimura et al., 2013, Katsuragi et al., 2015). p62 can form higher order structures like oligomers and helical filaments, and undergo liquid-liquid phase transition forming p62 bodies and p62 gels which are sites of autophagosome biogenesis and Nrf2 dependent antioxidant signaling (Yang et al., 2019, Kageyama et al., 2021, Komatsu, 2022).

Bacterial infections alter redox metabolism in the host cell and can activate Nrf2 dependent antioxidant signaling. Nrf2 activation can be protective for the host in some cases. For example, *Salmonella typhimurium* infection activates transcription of the Nrf2-target gene Ferroportin-1 which is a cellular iron exporter. This limits *Salmonella* access to iron, restraining its intracellular proliferation (Nairz et al., 2013). *Salmonella* infection also causes phosphorylation of p62 at S351 on the Salmonella containing vacuole, facilitating interaction with KEAP1 and activation of xenophagy (Ichimura et al., 2014). In infections with Uropathogenic Escherichia coli (UPEC), NRF2 activation in urothelial cells reduces ROS generation, inflammation, and promotes UPEC expulsion, therefore reducing the bacterial load. Pharmacologic activation of Nrf2 by sulforaphane causes resistance to *Pseudomonas aeruginosa* in mice (Harvey et al., 2011). Nrf2 signaling is also activated to control tissue damage for example in lung damage induced by *Staphyloccous aureus* infections (MacGarvey et al., 2012). Some bacteria like *Coxiella burnetti* exploit Nrf2 to promote intracellular growth. The pathogen stabilises p62, causing increase in its phosphorylation that activates Nrf2 signaling promoting bacterial growth in macrophages (Winchell et al., 2018). *Mycobacterium tuberculosis* infection also causes an upregulation of Nrf2-dependent gene expression thus promoting bacterial persistence in alveolar macrophages (Rothchild et al., 2019).

*Legionella pneumophila* infection has been associated with an increase in mitochondrial fragmentation and production of mitochondrial ROS, which was partially dependent on the bacterial effector protein MitF. We found phosphoribosyl-linked serine ubiquitination (PR-Ub) catalysed by the SidE effector proteins to be an important determinant of redox metabolism in infected cells. During infection, SdeA, SdeB, SdeC and SidE mediate phosphoribosyl-linked ubiquitination (PR-Ub) of the serine residues of substrate proteins (Qiu et al., 2016; Bhogaraju et al., 2016; Kotewicz et al., 2017). SdeA is structurally and mechanistically well-studied (Akturk et al., 2018; Dong et al., 2018; Kalayil et al., 2018). PR-Ub levels in infected cells are regulated by the action of PR-Ub specific deubiquitinases DupA and DupB (Shin et al., 2020). SdeA activity is fine-tuned by the Legionella glutamylase SidJ, which inhibits SdeA catalytic activity (Bhogaraju et al., 2019; Black et al., 2019; Gan et al., 2019). PR-Ub has been linked to fragmentation of the Rtn4-labeled tubular endoplasmic reticulum (ER), and to fragmentation of the Golgi body in Legionella-infected cells (Qiu et al., 2016; Kotewicz et al., 2017; Liu et al., 2021).

Proteomic analysis showed p62 to be a putative substrate of PR-Ub during Legionella infection. This suggested that autophagy may be regulated by PR-Ub of p62. However, *Legionella pneumophila* has a sophisticated mechanism to block the formation of autophagosomes using the bacterial effector protein RavZ, which irreversibly delipidates ATG8 proteins on autophagosomal membranes (Choy et al., 2012; Yang et al., 2017). This study shows how serine ubiquitination of p62 affects redox metabolism through its regulation of the p62-KEAP1-Nrf2 signaling axis. Using a combination of bacterial infection studies, biochemical experiments, and proteomics we have identified the PR-Ub site, and dissected the molecular mechanism by which serine ubiquitination of p62 controls KEAP1-mediated Nrf2 ubiquitination and degradation without affecting p62 phosphorylation. This mechanism results in a temporal regulation of redox metabolism in Legionella infected cells and leads to Nrf2 dependent remodelling of the host cell metabolome and proteome, which favour bacterial proliferation under oxidative stress.

## Results

### Mapping of serine ubiquitination sites in p62 during Legionella infection

Previous proteomic data revealed that p62 is PR-ubiquitinated in Legionella-infected cells (Shin et al., 2020). In order to validate the PR ubiquitination of endogenous p62 upon Legionella infection a GST pulldown assay was performed by incubating cell lysates from Legionella infected cells with the trapping mutant of DupA (GST-DupA(H67A)) followed by immunoblotting with antibodies against p62, GST and ubiquitin. ΔD (ΔDupA/B) Legionella infected cells had higher amounts of PR-Ub modified p62 compared to WT Legionella infected cells. Legionella strains lacking SidE enzymes (ΔS) are PR-Ub deficient (**Fig. 1A**). In addition, in order to identify the serine sites of p62 that are ubiquitinated we subjected purified maltose binding protein (MBP) tagged p62 to *in vitro* PR/ubiquitination reaction with SdeA, ubiquitin and NAD and the reaction products of this assay were subjected to mass spectrometry. High resolution electron-transfer dissociation (ETD) spectra of PR-ubiquitinated peptides identified serines S361 and S365 on p62 to be modified by PR-Ub (**Fig. 1B**). Based on the domain organization of the protein, it was noted that the identified PR-Ub sites of p62 were present in the unstructured loop between the KEAP1 interaction region (KIR) and the Ub-associated (UBA) domains (**Fig. 1B**). Mutation of three serine residues at S361, S365, S366 was necessary and sufficient to generate a PR-Ub deficient mutant of Myc-p62 (p62S361AS365AS366A) which was not PR-Ub modified upon bacterial infection (**Fig 1C**). His-tagged p62 peptide (comprising amino acids 321-440) was PR-Ub modified in an *in vitro* reaction, treated with or without GST-DupA, run on a SDS-PAGE and stained with Coomassie Brilliant Blue. *In vitro* PR-Ub resulted in 2 bands separated by ∼8KDa corresponding to p62(321-440) modified on one or different serines (p62-PR-Ub, p62-PR-Ub(2x)). Treatment with functional GST-DupA enzyme removed the PR-Ub from the p62 peptide proving that the modification on p62 was PR-Ub (**Fig. S1A**).

**Figure 1:**
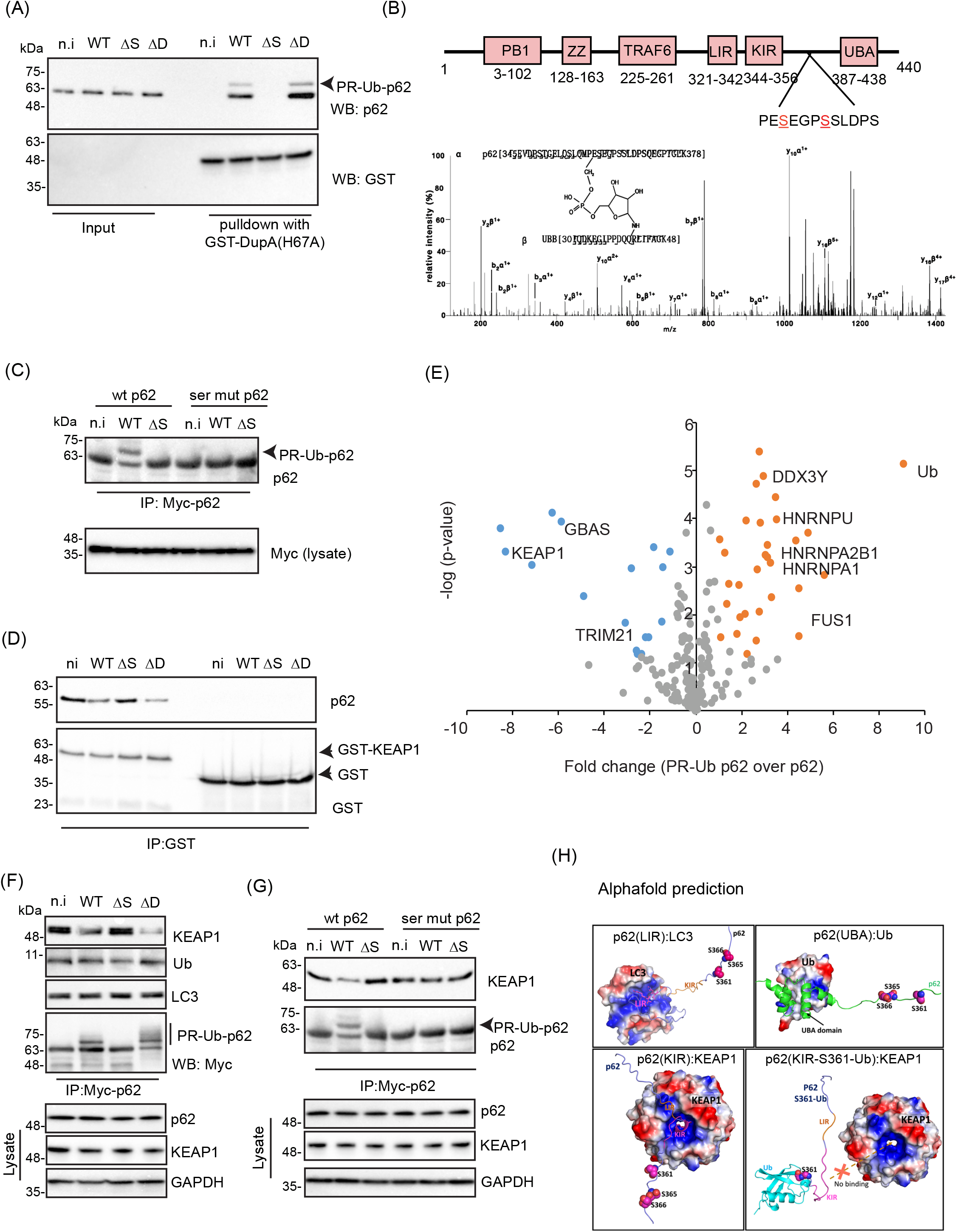
PR-Ub of p62 in Legionella infection reduces its interaction with KEAP1. **A.** HEK 293T cells expressing CD32 were infected with different strains of Legionella for 2 h. Lysates were used for GST pulldown with the DupA trapping mutant GST-DupA(H67A), followed by western blots with antibodies against p62, and GST. **B.** Domains of p62 and identification of site of PR-Ub **C.** HEK293T cells were transfected with myc tagged wild type or serine Ub deficient (S361AS365AS366Ap62) p62 and infected with Legionella for 2h followed immunoprecipitation with Myc resin followed by western blotting **D.** Lysates from cells infected with Legionella for 2 h were incubated with GST tagged KEAP1 or GST alone in a GST pulldown assay, followed by immunoblotting with GST and p62 antibodies **E.** GST-p62(amino acids 321-400) of p62 purified from E. coli, was PR-ubiquitinated it in an *in vitro* reaction or left unmodified and incubated it with cell lysate from HEK293T cells infected with WT Legionella in a GST pull down assay, followed by mass spectrometric identification of interactors. **F.** A549 cells were transfected with Myc-p62 and infected with indicated Legionella strains for 2h, followed by immunoprecipitation of myc-p62 and immunoblotting with LC3, Ub, KEAP1, and Myc antibodies. **G.** A549 cells were transfected with myc tagged wild type p62 or (S361AS365AS366A)p62 and infected with indicated Legionella strains for 2h, followed by immunoprecipitation of myc-p62 and immunoblotting with KEAP1, and p62 antibodies. **H.** Alpha fold prediction of the complex formed between the p62 peptide (amino acids 321-400) and LC3 or KEAP1 (n.i-not infected, WT-wild-type Legionella, ΔS-ΔSidE Legionella, ΔD-ΔDupA/B Legionella)

### PR-Ub of p62 inhibits its interaction with KEAP1

Since the identified PR-Ub site was in close proximity of the LIR, KIR and UBA domains of p62 we checked whether interaction of p62 with any of these interactors was affected by its PR-Ub. GST-tagged ubiquitin and LC3 interacted with PR-Ub modified p62 (**Fig. S1B**), but GST-KEAP1 showed reduced interaction with PR-Ub p62 in WT and ΔD Legionella infected lysates (**Fig 1D**). Next, we purified a peptide spanning amino acids 321-400 of p62 from E. coli, PR-ubiquitinated it in an *in vitro* reaction and incubated it with cell lysate from HEK293T cells infected with Legionella in a GST pull down assay, followed by mass spectrometric identification of interactors. Upon PR-ubiquitination, p62 (321-400) bound significantly less KEAP1 supporting the data in Fig 1D. We also saw a significant enrichment of several proteins involved with RNA metabolism, HDACs and DNA damage related proteins which preferentially bound to PR-Ub modified p62 peptide (**Fig. 1E and S1C**), which will be validated by utilising the full-length protein in a follow up study.

To test for the interaction between p62 with KEAP1 in cells we used Myc-p62 expressing A549 cells infected with wild type and mutant Legionella strains for 2h to enrich modified myc-p62 and checked for co-precipitation with KEAP1. PR-Ub modified p62 present in WT and ΔD infected cells bound to less KEAP1when compared to uninfected or ΔS infected cells (**Fig. 1F**). This difference in p62-KEAP1 interaction between WT and ΔS *Legionella* infection was insignificant when the PR-Ub deficient mutant of p62 (p62S361AS365AS366A) was used (**Fig. 1G**). In order to understand the structural basis of how serine ubiquitination of p62 alters binding to its interaction partners we performed Alphafold-based complex prediction of LIR, KIR and UBA domains of p62 with LC3, KEAP1 and ubiquitin, respectively. As expected, the complexes predicted were well in agreement with already existing crystal structures of LIR with LC3 (PDB Id: 2ZJD) and KIR with KEAP1 (PDB ID: 3WDZ), however the structure of p62-UBA domain in complex with ubiquitin is unknown. These complexes clearly show that the site of PR-ubiquitination (S361/365/366) are in very close proximity of KIR domain but not LIR and UBA domain. We further added ubiquitin on Serine 361 of p62 and predicted the complexes. As expected, the binding of Ser361 ubiquitinated p62 lost binding to KEAP1, this largely due to the steric exclusion of KIR from the KEAP1 pocket due to presence of ubiquitin (**Fig. 1H**).

### PR-Ub of p62 inhibits recruitment of KEAP1 to intracellular bacteria

Previous reports showed that ∼10% Legionella containing vacuoles were positive for p62 in the first 2-3 h post infection (Omotade and Roy, 2020). To check if p62 was recruited to intracellular Legionella upon infection and whether this was affected by PR-Ub we checked the localization of WT and PR-ub deficient myc-p62 2 h.p.i. We observed about 40% of infected A549 cells expressing Myc-p62 to have at least one intracellular bacterium which was positive for p62. The PR-Ub deficient mutant of p62 (p62S361AS365AS366A) was also bound to intracellular bacteria in a similar manner (**Fig. 2A**). This suggest that p62 is recruited to intracellular bacteria vacuoles in the early part of the infection cycle and this is independent of its PR-Ub. Similarly, previous studies in *Salmonella* have shown the presence of p62 and KEAP1 at intracellular bacterial vacuoles. Phosphorylation of p62 at S351 was important for p62-KEAP1 interaction and recruitment of KEAP1 to p62 positive bacterial vacuoles (Ishimura et al., 2014). To test whether endogenous KEAP1 was recruited to intracellular Legionella, A549 were depleted of endogenous p62 by treatment with p62 siRNA for 48h, followed by reconstitution with WT and PR-Ub deficient Myc-p62. Cells were then infected with WT or ΔS Legionella for 2 h followed by immunofluorescence analysis of infected cells. This experiment showed that PR-Ub of p62 prevented KEAP1 recruitment to intracellular bacteria. KEAP1 was recruited to bacteria in ΔS Legionella infection and in cells expressing PR-Ub deficient p62 (**Fig. 2B**). A549 cells depleted of endogenous p62 followed by reconstitution with Myc tagged WT or PR-Ub deficient p62 were infected with WT Legionella. Lysates were collected from infected cells at different time points post infection and analysed for phosphorylation of p62 at S351. Legionella infection triggered phosphorylation of p62 at S351; the amount of p-S351-p62 increased with time and was unaffected by PR-Ub. The p62-KEAP1 interaction was reduced by PR-Ub at 1 h.p.i and at 2 h.p.i. At later time-points, PR-Ub modified p62 was not detected and the p62-KEAP1 interaction was similar to that in uninfected cells (**Fig. 2C**).

**Figure 2:**
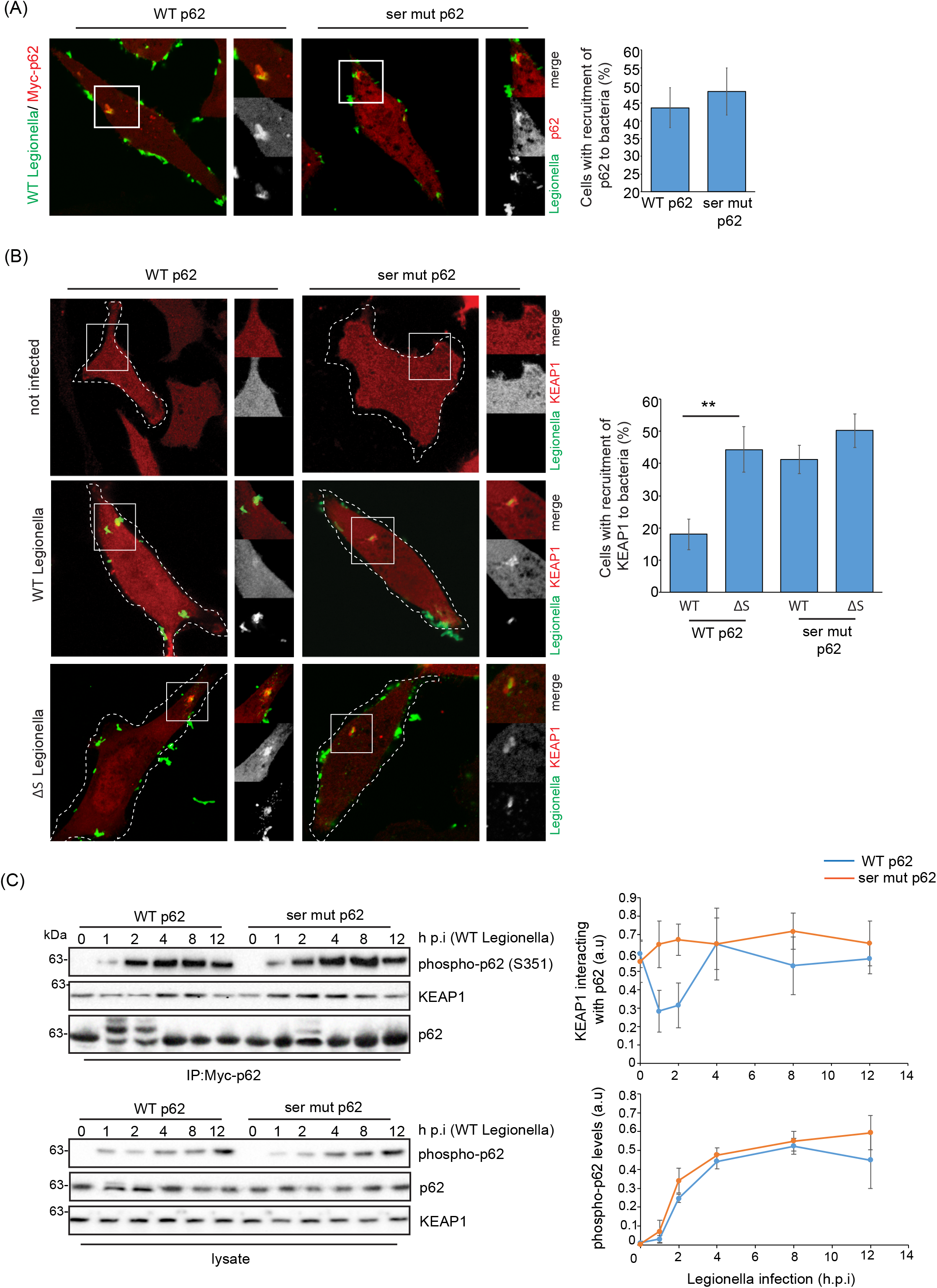
KEAP1 recruitment to intracellular bacteria is blocked by PR-Ub of p62. A. A549 cells expressing myc tagged wild type p62 or S361AS365AS366Ap62) p62 B. were infected with WT Legionella for 2h followed by fixation and immunostaining to visualize recruitment of myc-p62 to bacteria. 60 cells were analyzed inFIJI manually to check recruitment of p62 to bacteria and numbers were plotted in MS Excel. C. A549 cells expressing myc tagged wild type p62 or S361AS365AS366Ap62) p62 D. were infected with WT or ΔS Legionella for 2h followed by fixation and immunostaining with KEAP1, Legionella antibodies to visualize recruitment of KEAP1 to bacteria. 60 cells were analyzed in FIJI manually to check recruitment of KEAP1 to bacteria and numbers were plotted in MS Excel. Error bars, standard deviation, ** 0.01 >p≥0. 001. E. A549 cells expressing myc tagged wild type p62 or (S361AS365AS366Ap62) p62 F. were infected with WT Legionella and lysed at indicated time-points followed by immunoprecipitation of p62-myc and immunoblotting with antibodies against p62, P(S351)-p62, and KEAP1.Graphs indicate quantitation of band intensities of 3 immunoblots taken from 3 independent experiments. Error bars indicate standard deviation, ** 0.01 >p≥0. 001.

### PR-Ub of p62 modulates Nrf2 levels in a KEAP1 dependent manner

Next, we tested how Nrf2 levels are regulated during the infection time course by using A549 cells depleted of endogenous p62 by siRNA treatment and infected with WT Legionella, lysed at different time-points post infection and analysed by immunoblotting to detect Nrf2 levels. Nrf2 levels were slightly higher in control siRNA treated cells compared to cells treated with p62 siRNA. Nrf2 levels increased with the progress of the infection suggesting an activation of Nrf2 signaling at later stages of infection (4-12 h.p.i). Reconstitution of p62 depleted cells with WT or PR-Ub deficient p62 followed by bacterial infection showed a dependence of Nrf2 levels on PR-Ub of p62 at 4 h.p.i and 6 h.p.i. At these time-points cells expressing the PR-Ub deficient p62 had higher levels of Nrf2 than those with WT-p62. This is consistent with previous findings that upon PR-Ub, p62 did not bind to KEAP1 (**Fig. 1D-G**). This caused KEAP1 to interact with Nrf2 and cause its ubiquitination and proteasomal degradation (**Fig. 3A**). A549 cells depleted of endogenous KEAP1 with siRNA, infected with either WT or ΔS Legionella had higher Nrf2 levels throughout the time course of infection (**Fig. 3B**). This proved that both p62 and KEAP1 are important in regulating Nrf2 levels during the course of Legionella infection.

**Figure 3:**
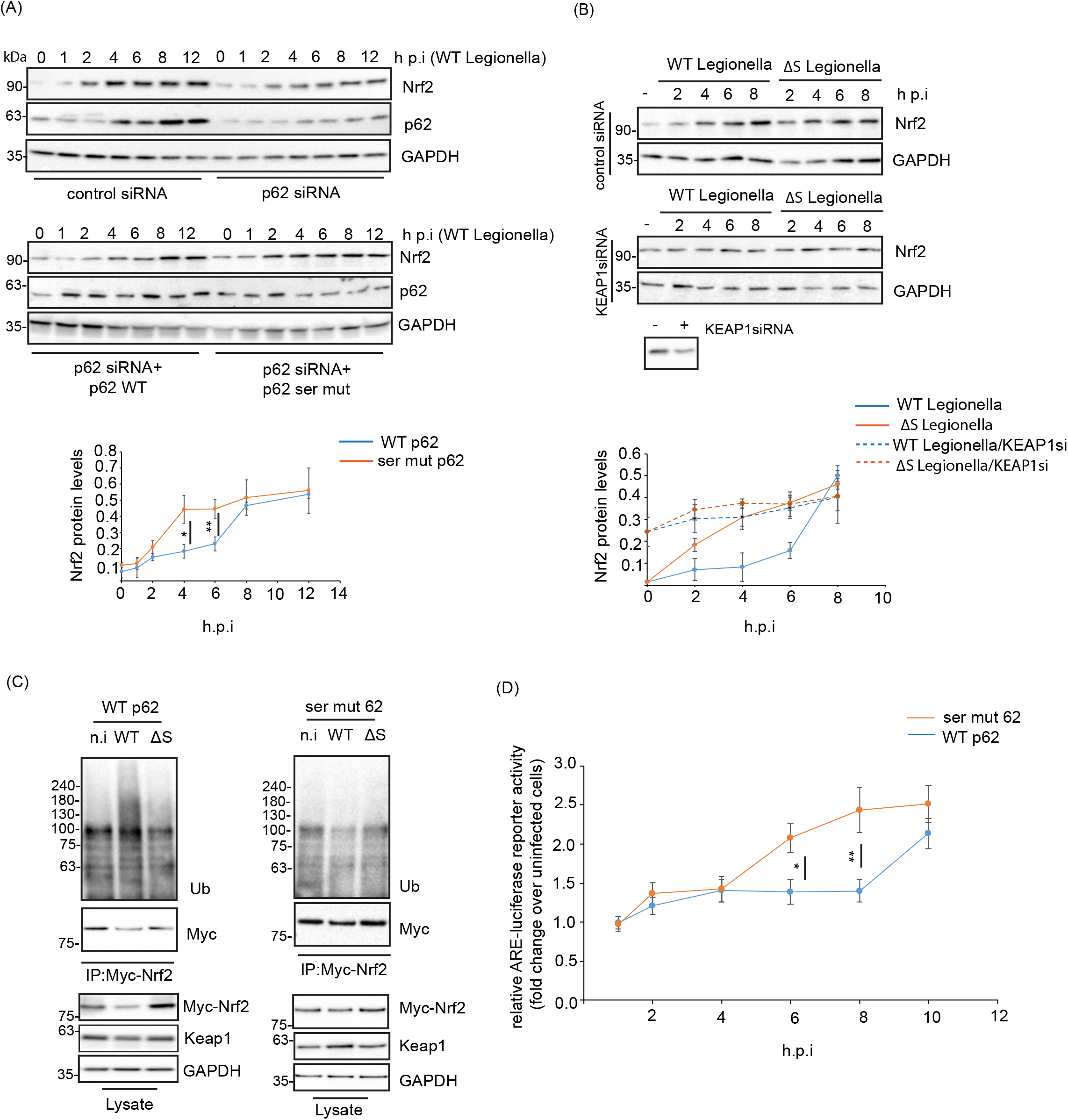
PR-Ub of p62 promotes Nrf2 ubiquitination and degradation. **A.** A549 cells were treated with p62 siRNA for 48h followed by reconstitution with wild type or PR-Ub deficient (S361AS365AS366A) p62 for 24h. these are then infected with WT Legionella and lysed at indicated time-points. Lysates are blotted with p62, Nrf2, GAPDH antibodies. Graphs indicate quantitation of band intensities of 3 immunoblots taken from 3 independent experiments. Error bars indicate standard deviation. **B.** A549 cells were treated with KEAP1 siRNA for 48h followed by infection with WT or ΔS Legionella and lysed at indicated time-points. Lysates are blotted with Nrf2, and GAPDH antibodies. Graphs indicate quantitation of band intensities of 3 immunoblots taken from 3 independent experiments. Error bars indicate standard deviation. **C.** A549 cells expressing Myc-Nrf2 and p62 (wild type or PR-Ub deficient) were infected with WT or ΔS Legionella for 6h, Nrf2 ubiquitination was probed by immunoprecipitating Nrf2-Myc and immunoblotting with Ub. **D.** A549 cells cotransfected with ARE (Nrf2 promoter)-luciferase reporter construct and wild type or PR-Ub deficient p62 were infected with Legionella for indicated times followed by measurement of luciferase reporter activity. Graph represents ARE-luciferase activity expressed as the fold change from uninfected cells. Error bars indicate standard deviation, * 0.05>p≥0.01, ** 0.01>p≥0.001.

p62 depleted cells were co-transfected with WT or PR-Ub deficient p62 and Myc-Nrf2 followed by infection with WT or ΔS Legionella for 6 h. Nrf2 was enriched from cell lysates on Myc resin and its ubiquitination was assayed by immunoblotting with anti-ubiquitin antibody. A greater amount of ubiquitinated Nrf2 was observed in WT Legionella infected cells compared to uninfected or ΔS infected cells. This difference of Nrf2 ubiquitination was absent in cells expressing PR-Ub deficient p62 highlighting the importance of p62 PR-Ub in regulation of Nrf2 ubiquitination and degradation (**Fig. 3C**).

### Nrf2-mediated antioxidant responses are dependent on PR-Ub of p62

Myc-Nrf2 expressing A549 cells were infected with different strains for 6 h followed by fractionation of lysates into nuclear and cytosolic fractions. WT and ΔD Legionella infected cells had significantly lower levels of Myc-Nrf2 in nuclear fractions suggesting lower Nrf2 dependent transcription (**Fig. S1D**). To assay Nrf2 dependent transcription from antioxidant response elements we used A549 cells expressing the reporter ARE-Luciferase. These cells were depleted of endogenous p62 by treatment with p62 siRNA followed by reconstitution with WT or PR-Ub deficient p62, infected with WT Legionella and assayed for ARE-luciferase activity at different time points of infection. Cells expressing WT p62 had lower Nrf2 dependent ARE transcription at 6 and 8 h.p.i, when compared to cells expressing mutant p62 (**Fig. 3D**). At 6 h.p.i WT Legionella infected cells had lower Nrf2 dependent transcription than ΔS Legionella infected cells (**Fig. S1E**).

### Downregulation of Nrf2 dependent genes upon acute Legionella infection

A549 cells were infected with WT or ΔS Legionella for 6 h followed by extraction of total mRNA. Next generation RNA sequencing was performed using the Illumina platform to check differential expression of genes. Expression of several known Nrf2 target genes including Hmox1 (heme oxygenase 1), MT1X (Metallothionein 1X), TXNRD1 (Thioredoxin Reductase 1), SOD2 (superoxide dismutase 2), CAT (catalase), SQSTM1 (sequesterome1/p62), and SLC48A1 (Solute Carrier Family 48 Member 1), NQO1 (NADPH dehydrogenase quinone 1) are higher in ΔS Legionella infection compared to cells infected with WT Legionella (**Fig. 4A and S2A**).Gene Ontology (GO) analysis of differentially expressed genes using the online server MetaScape identified oxidative stress response, peroxisome and pyruvate metabolism and TCA cycle to be the pathways upregulated in ΔS Legionella infection when compared to infection with the wild-type bacteria. WT Legionella infection led to upregulation of genes related to other GO classes like VEGFR signaling, DNA damage response and cytokine signaling (**Fig S2B and S2C**). Nrf2 (also known as NFEL2) mediated regulation of gene expression was identified as the most transcriptionally upregulated pathway in ΔS Legionella infection. Transcriptional pathways regulated by NFKB, p53 and HIFα are upregulated in WT Legionella infection (**Fig. S3A and S3B**).

**Figure 4:**
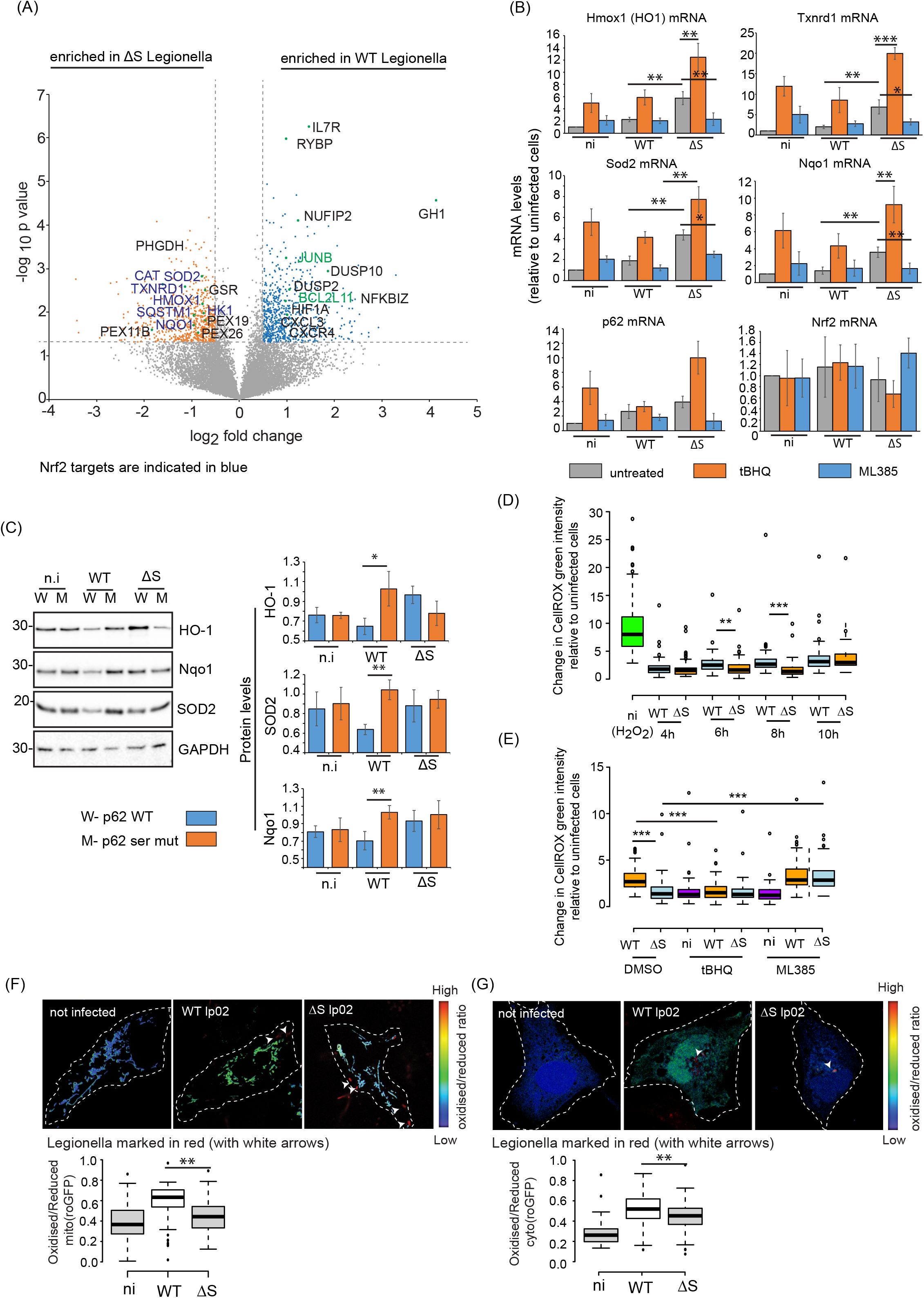
PR-Ub of p62 regulates Nrf2 dependent gene expression. A. Differential expression of genes between WT and ΔS Legionella infection B. RT-PCR to measure expression of Nrf2 targets in A549 cells infected with WT or ΔS Legionella for 6 h in presence of 10µM tBHQ or ML385. C. A549 cells expressing wild type or PR-Ub deficient p62 were infected with WT or ΔS Legionella for 6 h followed by immunoblotting of lysates with indicated antibodies. Graphs represent band intensities from 3 experiments, error bars indicate standard deviation. * 0.05>p≥0.01, ** 0.01>p≥0.001. D. Total ROS measured in infected cells at indicated time-points by oxidative stress indicator CellROX green. E. CellROX green measurements at 8 h.p.i in cells infected with Legionella in presence of 10µM tBHQ or ML385. F. Mitochondrial roGFP measurements in Legionella infected cells 8 h.p.i, ** 0.01>p≥0.001. G. Cytosolic roGFP measurements in Legionella infected cells 8 h.p.i, ** 0.01>p≥0.001.

Next, we infected A549 cells with WT or ΔS Legionella in the presence of the Nrf2 activator tBHQ (tertiary butylhydroquinone) or the Nrf2 inhibitor ML385 for 6 h followed by extraction of mRNA and quantitative PCR to measure the differential expression of Nrf2 target genes. Similar to our RNA sequencing data, WT Legionella infected cells had lower levels of Nrf2 target genes including HMOX1, TXNRD1, SOD2, NQO1 and SQSTM1 when compared to cells infected with ΔS Legionella. Upon treatment with tBHQ WT Legionella infected cells (where Nrf2 is degraded by KEAP1) show a slight to moderate increase in expression of Nrf2 targets, while ΔS Legionella infected cells show a significant increase in expression of Nrf2 target genes. On the other hand, treatment with ML385 had almost no effect in expression levels of Nrf2 targets in WT Legionella infected cells as Nrf2 mediated gene expression is already blocked through PR-Ub mediated regulation of the p62-KEAP1-Nrf2 signaling pathway. ΔS Legionella infected cells treated with ML385 show a significant reduction in Nrf2 dependent gene expression (**Fig. 4B**). We also tested protein levels of Nrf2 regulated antioxidants SOD2, HO-1, and NQO1 in Legionella infected cells. Expression of these antioxidants were lower in WT Legionella infection when compared to ΔS infection. Expression of the PR-Ub deficient mutant of p62 (p62S361AS365AS366A) in p62 depleted cells abolished this difference in antioxidant levels (**Fig. 4C**). Collectively these experiments indicated that in WT Legionella infection, Nrf2 dependent expression of antioxidants is blocked through PR-Ub of p62; KEAP1 can interact with Nrf2 causing its ubiquitination and proteasomal degradation. In absence of PR-Ub (as in ΔS Legionella infection), phosphorylated p62 (pS351-p62) binds to KEAP1, leading to stabilization of Nrf2 and expression of its downstream targets.

### Cellular ROS levels are regulated in a PR-Ub dependent manner

Since PR-Ub of p62 regulates antioxidant levels, we decided to check the levels of total cellular ROS at different time-points of infection. For this Legionella infected cells were loaded with the oxidative stress sensitive fluorogenic probe CellROX green for 30min, fixed with paraformaldehyde and imaged by high content fluorescence microscopy. WT Legionella infection led to higher ROS levels at 6 and 8 h.p.i when compared to infection with the ΔS strain. At 10 h.p.i ROS levels were similar between the WT and ΔS infected cells (**Fig. 4D**). Treatment of infected cells with tBHQ and ML385 reduced and increased ROS levels respectively underlining the dependence of cellular ROS in infected cells on the Nrf2 pathway (**Fig. 4E**).

A549 cells were transfected with a mitochondria targeted redox-sensitive GFP probe (mito-roGFP) which is a ratiometric reporter of oxidative stress in live cells. Similar to the total ROS measurements, mitochondrial ROS was higher in WT Legionella infection compared to the ΔS bacteria at 8 h.p.i (**Fig. 4F**). Infection of cells containing a cytosolic variant of redox GFP also showed similar results (**Fig. 4G**).

### PR-Ub of p62 remodels the proteome in a Nrf2 dependent manner

We infected A549 cells with WT or ΔS Legionella in the presence of tBHQ or ML385 for 8 h followed by protein extraction and analysis of the whole cell proteome and phosphoproteome by mass spectrometry to identify the Nrf2 dependent changes in the proteome (**Fig. 5A**). In uninfected cells treatment with tBHQ for 8 h lead to the significant increase in expression of known Nrf2 dependent genes including SQSTM1, antioxidants (NQO1, TXNRD1, GPX2, SOD2, CAT), and glycolysis enzymes (HK1, G6PD, TALDO1). On the other hand, treatment with ML385 led to an increase in enzymes related to fatty acid metabolism (ACOX1, DGAT1, HACD2, ACADS, ACSM5, SCD, SELENOI), proteins related to mitochondrial respiration (MT-CO3, MT-CO1, SDHD, ATP6, COX7C), and proteins related to steroid biosynthesis (EBP, HMGCR, CYP24A1, SDHC, APOC3, MSMO1, CYP5A1) (**Fig. 5B and 5C**). This indicated a Nrf2 dependent remodelling of the proteome in A549 cells. Upon infection of cells with WT Legionella, both tBHQ and ML385 did not have any significant impact on the change in Nrf2 dependent antioxidants which was consistent with the high ROS levels observed in WT Legionella infection. Though these cells were less responsive to Nrf2 activation, other pathways related to metabolic changes and mitochondrial respiration were altered. tBHQ treatment increased levels of mitochondrial proteins (COX7C, NDUFB8, NDUFC1, ATP5MG, ATP5MF, UQCRQ, NIPSNAP1), enzymes involved in glucoronate metabolism (UGT1A1, UGT2A3, UGT2B7, CYP1B1, CYP3A5) (**Fig. S4A-D**). Infection of cells with ΔS Legionella led to an increase in Nrf2 dependent antioxidants and metabolic enzymes. Blocking Nrf2 dependent gene expression with ML385 led to increased expression of peroxisomal genes (PEX2, PEX13, PMP34, MPV17, HSD17B), and other genes related to detoxification of ROS (TXN2, TXNDC1, TMX4, TMX1, PRDX3) (**Fig. 5D, 5E, S4E**). Analysis of the transcription regulatory networks regulated in a tBHQ dependent manner in uninfected, WT and ΔS Legionella infection utilising the online tool TRRUST (transcriptional regulatory <u>r</u>elationships <u>u</u>nravelled by <u>s</u>entence-based text-mining, Han et al, 2015) identified NRF2 to be within the top hits altered by tBHQ treatment in uninfected and ΔS Legionella infected cells; WT Legionella infection did not show induction of Nrf2 dependent genes (**Fig. S5A-C**). Phosphoproteome analysis of Legionella infected cells treated similarly as shown in Fig. 5A led to identification of 1525 phosphosites including serine, threonine and tyrosine residues (**Fig. 5F and 5G**). 67 proteins and 164 proteins were differentially phosphorylated in WT and ΔS Legionella infection respectively. GO analysis of proteins differentially regulated by phosphorylation in WT Legionella infection led to enrichment of terms related to cellular metabolism (**Fig. 5H**). Proteins differentially regulated by phosphorylation in ΔS Legionella infection included those related to RNA binding, transcription and cellular response to stress (**Fig. S5D**).

**Figure 5:**
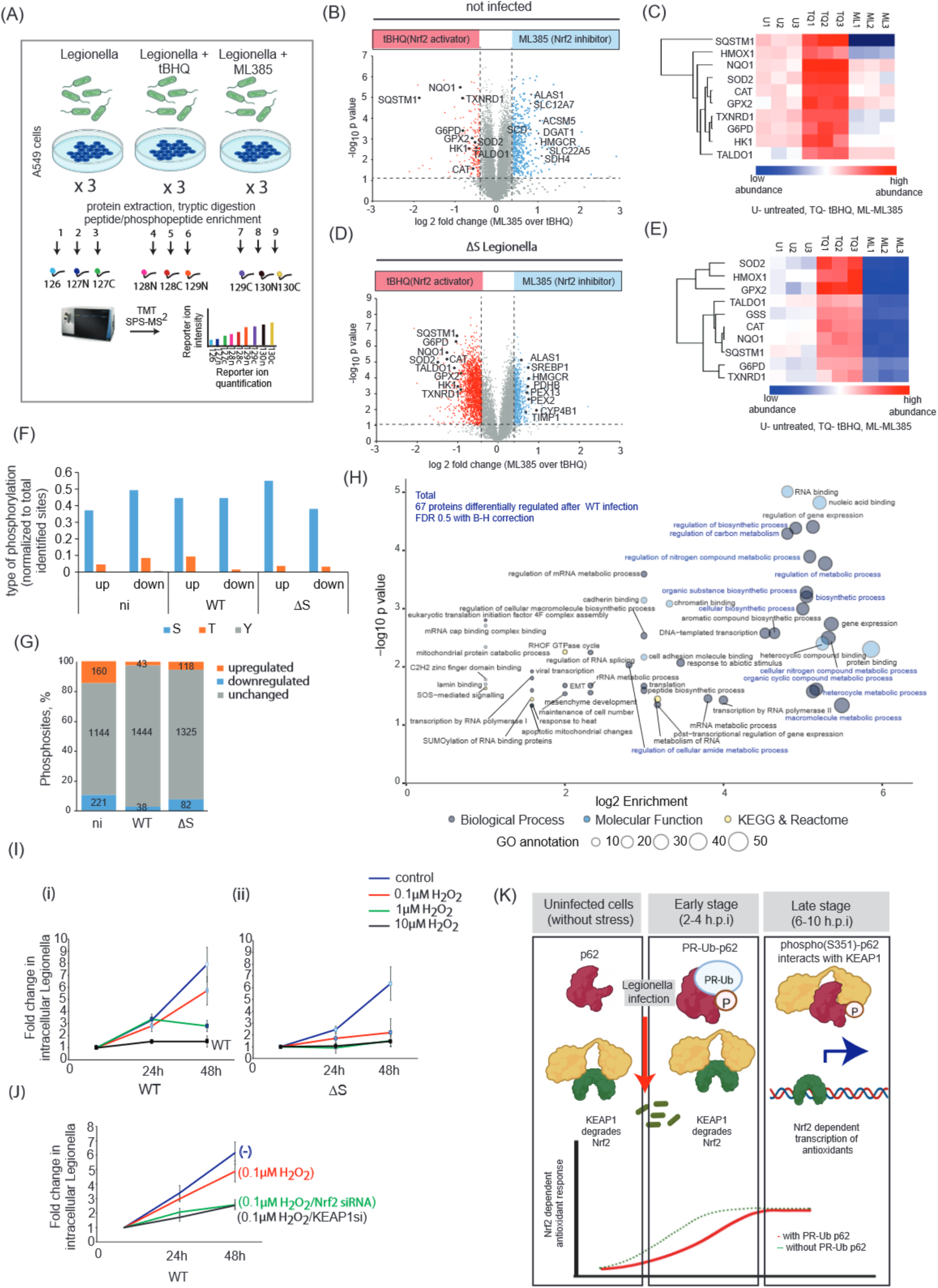
Legionella infection remodels the host proteome in a Nrf2 dependent manner. A. Schematic of experimental strategy B. Volcano plot showing changes in the proteome of A549 cells when treated with 10µM tBHQ or 10µM ML385 for 8 h. C. Levels of Nrf2 dependent genes in the experiment indicated in panel b. D. Volcano plot showing changes in the proteome of A549 cells infected with ΔS Legionella in presence of 10µM tBHQ or 10µM ML385 for 8 h. E. Heatmap showing levels of Nrf2 pathway proteins in the experiment indicated in panel d. F. Proportion of S, T, Y phosphosites identified upon phosphoproteome analysis of Legionella infected cells at 8 h.p.i. G. Differentially regulated phosphosites in uninfected, WT, ΔS Legionella infection H. GO analysis of the 67 proteins differentially regulated by phosphorylation during WT Legionella infection. Metabolism related processes are marked in blue. I. A549 cells were infected with WT or ΔS Legionella for 12 h, 24 h or 48 h in presence of H2O2 followed by estimation of intracellular bacterial load measured by a bacterial replication assay. J. A549 cells depleted of endogenous KEAP1 or Nrf2 were infected with Legionella for 12 h, 24 h or 48 h in presence of H2O2 followed by estimation of intracellular bacterial load measured by a bacterial replication assay. K. Summary showing effect of PR-Ub on the p62-KEAP1-Nrf2 pathway.

### PR-Ub mediated remodelling of oxidative metabolism favours bacterial replication in host cells

To check how the redox modulation of the transcriptome and proteome during infection affects bacterial proliferation in the host cell we performed an intracellular replication assay of Legionella under conditions of oxidative stress induced by increasing concentration of H_2_O_2_. We found that replication of WT bacteria was unaffected by a low concentration of H_2_0_2_ (0.1µM), However, the same concentration of H_2_0_2_ was enough to cause a significant reduction in the intracellular replication of ΔS Legionella. Higher concentrations of H_2_0_2_ (1 µM and 5 µM) reduced bacterial replication of both WT and ΔS Legionella (**Fig. 5I**). 5 µM H_2_0_2_ also reduced the viability of A549 cells in an MTT assay (**Fig. S5E**). Depletion of either Nrf2 or KEAP1 in host cells reduced the intracellular replication of WT Legionella in presence of 0.1µM H_2_0_2_ (**Fig. 5J**).

Collectively, this study shows how serine ubiquitination of p62 prevents Nrf2 activation in the first few hours of infection. By 4 h.p.i, PR-Ub-p62 levels drop significantly, allowing transcriptional activation of Nrf2 target genes at a later stage of infection (**Fig. 5K**).

## Discussion

This study shows how Legionella utilises serine ubiquitination to fine-tune redox metabolism of the infected cell through the regulation of the p62-KEAP1-Nrf2 signaling axis. Upon PR-Ub, p62 is sterically prevented in binding to KEAP1, which is then available to interact with Nrf2, causing its ubiquitination and proteasomal degradation. This mechanism prevents the premature activation of the Nrf2 dependent redox response in the early stage of infection. The activity of DUPs causes the levels of PR-Ub-p62 to drop and completely disappear by 4 h.p.i., (Shin et al., 2020); removing the p62-KEAP1 mediated inhibition on Nrf2 activation, providing a temporal regulation of the transcription of Nrf2 target genes

In parallel, Legionella infection triggers the phosphorylation of p62 at S351 within the first 1h of infection. In absence of PR-Ub (as in ΔS Legionella), pS351-p62 competes with Nrf2 for binding to KEAP1. As a result, Nrf2 is stabilised and transcription of Nrf2 targets are activated at an early phase of infection. Therefore, PR-Ub does not interfere with the phosphorylation of p62 but delays the activation of Nrf2 to a later stage of infection.

*Legionella pneumophila* has several effectors which block autophagy in the host cell. The most prominent of these bacterial proteins is RavZ which delipidates ATG8 proteins, blocking the formation of autophagosomes. Therefore, Legionella infected cells can be considered to be an autophagy deficient system where there may be build-up of ubiquitinated autophagic cargo (like ubiquitinated bacteria, defunct mitochondria, or protein aggregates) which can bind p62, form p62 oligomers and lead to activation of the p62-KEAP1-Nrf2 signaling cascade. In the absence of a functional autophagy-lysosomal system, phospho(S351)-p62 remains elevated throughout the infection time-course (measured up to 12 h.p.i). Though p62 positive autophagosomes are not formed during infection, our studies and others have reported the recruitment of p62 to ubiquitinated Legionella (Omotade and Roy, 2020). From the immunofluorescence analysis, we infer that serine ubiquitination of p62 does not affect its recruitment to the bacterial phagosome in the early phase of infection but prevented the recruitment of KEAP1 to the bacterial phagosomes. In absence of PR-Ub, the p351-p62-KEAP1 interaction on ubiquitinated bacterial phagosomes lead to the stabilisation and nuclear translocation of Nrf2.

Nrf2 is an important signaling hub which regulates several aspects of metabolism. Our results indicate that Nrf2 serine ubiquitination has an impact on transcription of ROS detoxification enzymes (like TXNRD1, NQO1, SOD2, etc) and their regulation on the cellular ROS levels at different time-points of infection. It is also important to consider that Nrf2 target genes include several metabolic enzymes including those involved in glycolysis and pyruvate metabolism. These include HK1(hexokinase1), TALDO1 (transaldolase1), G6PD1(glucose-6-phosphate dehydrogenase), PGD (phosphogluconate dehydrogenase). Our RNAseq data showed that these Nrf2 target genes were also downregulated in WT Legionella infection suggesting a differential regulation of carbon flux through glycolysis and the pentose phosphate pathway. Pathway analysis to compare gene expression differences between WT and ΔS Legionella infection identified carbon metabolism, carbohydrate metabolism, pyruvate metabolism, and TCA cycle to be within the top 20 hits suggesting a serine ubiquitin dependent remodeling of metabolism, which may be partially dependent on Nrf2. Previous studies on macrophages infected by Legionella have reported changes in mitochondrial respiration and glycolysis induced by bacterial effectors. A shift to glycolytic (Warburg-like) metabolism at a later phase of infection promotes proliferation of Legionella (Escoll et al., 2017). Our studies suggest that Nrf2 dependent regulation of gene expression may have an important role in determining the metabolic switch to glycolysis. Nrf2 signaling is also linked with inflammation; cross-talk of Nrf2 signaling with the NFκB pathway has been characterized in several cases (Cuadrado et al., 2014, Brazao et al., 2022). The Nrf2 target gene HMOX1 encodes heme oxygenase 1 which modulates intracellular iron metabolism and negatively regulates inflammation (Wu et al., 2021). Whether and how iron metabolism and inflammation are regulated by serine ubiquitination will be further studied in the future.

It is interesting to contemplate how the serine ubiquitination mediated regulation of the redox response affects intracellular Legionella. The intracellular replication assay suggests that WT Legionella can survive and proliferate in cells treated with the oxidative stressor (1µM H202), while in the case of ΔS Legionella, the intracellular replication is significantly compromised. The Nrf2 dependent remodelling of gene expression contributes to the better tolerance of WT bacteria to oxidative stress as siRNA mediated depletion of Nrf2 and KEAP1 reduced bacterial replication significantly. The better tolerance to oxidative stress may be due to differential regulation of metabolic pathways that affect nutrient availability to the bacteria It may also be affected by differences in activation of cell death pathways between WT and ΔS Legionella infection. Studies on uropathogenic Ecoli (UPEC) have demonstrated how Nrf2 dependent transcription of Rab27B helps in expulsion of bacteria from urothelial cells at a late stage of infection (Joshi et al., 2021). It may be possible to envision circumstances where the optimal timing of bacterial release from Legionella infected cells is influenced by PR-Ub dependent Nrf2 activation, which in turn would influence the intracellular load of bacteria at a given time.

## Materials and methods

### Plasmid construction

**cDNA for human p62 (NCBI reference number:** NM_003900.5) **was cloned with a N terminal Myc tag in pCMV3 vector from SinoBiological (Catalog no:** Cat: CV011) and in pcDNA3-HA2-Keap1 and pCDNA3-Myc3-Nrf2 are gifts from Yue Xiong (Addgene plasmid # 21556; and # 21555; respectively) (Furukawa et al., 2005). Matrix-roGFP and cyto-roGFP was a gift from Paul Schumacker (Addgene plasmid # 49437 and Addgene plasmid # 49435) (Waypa et al., 2005). pLminP_Luc2P_RE4 was a gift from Ramnik Xavier (Addgene plasmid # 90338), (O’Connell et al., 2016). GST tagged KEAP1 and Ubiqutin were generated by cloning human cDNA sequences into pGEX6P1. Point mutations in p62 were introduced by standard site directed mutagenesis.

### Protein purification

GST-tagged DupA, KEAP1 and Ubiquitin were purified from *E. coli* as previously described (Shin et al., 2020). Briefly, BL21(DE3) competent cells (NEB) were transformed with plasmids and grown in lysogeny broth at 37 °C to an OD_600_ of 0.6–0.8. Protein expression was induced by adding 0.5 mM isopropyl-D-thiogalactopyranoside (IPTG) overnight at 18 °C. Lysates were incubated with glutathione Sepharose resin pre-equilibrated with lysis buffer (50 mM Tris-HCl (pH 7.5), 150 mM NaCl, 3mM DTT), and non-specific proteins were cleared by washing twice with wash buffer (50 mM Tris-HCl (pH 7.5), 300 mM NaCl, 3mM DTT. Proteins were eluted in 50 mM Tris-HCl (pH 7.5), 150 mM NaCl, 15 mM glutathione and exchanged into storage buffer (20 mM Tris-HCl pH 7.5, 100 mM NaCl) before further use.

### Antibodies and siRNA

Human siRNA was used from Santa Cruz Biotechnology. P62/SQSTM1 siRNA (sc-29679). Keap1 siRNA (h) (sc-43878), Nrf2 siRNA (h) (sc-37030).

We used the following antibodies and dilutions: p62 (cat. no. 610832, BD Biosciences, 1:1000), KEAP1 (Cell Signaling Technology, cat no #4678, 1:1000), Nrf2 antibody (Santa Cruz biotechnology, sc-365949; 1:1000), GAPDH (cat. no. 2118, Cell Signaling Technology; 1:2000), Myc beads (cat. no. sc-40 AC, Santa Cruz Biotechnology), LC3 (cat. no. 2775, Cell Signaling Technology; 1:2000), phospho-p62 (cat. no. 95697, Cell Signaling Technology; 1:1000), Ubiquitin (Cat. no: 3933, Cell Signaling Technology, 1:1000), HO1 (cat no: sc-136960, Santa Cruz Biotechnology, 1:2000), NQO1 (ab34173, abcam, 1:2000), SOD2 (ab13533, abcam, 1:1000), GST resin (cat no: 16101, Pierce, Thermofisher Scientific), Legionella antibody (ab

### Immunoprecipitation and western blotting

Cells were lysed in 50 mM Tris-HCl (pH 7.5) containing 150 mM NaCl and 1% Triton X-100. For the immunoprecipitation of Myc-p62 or Myc-Nrf2, lysates containing 1 mg protein were incubated with myc beads in immunoprecipitation buffer (50 mM Tris-HCl pH 7.5, 150 mM NaCl, 0.5% Triton X-100), then washed (50 mM Tris-HCl pH 7.5, 300 mM NaCl, 0.5% Triton X-100) and boiled with SDS sample buffer. For western blotting, 20-µg samples were loaded onto 10% Tris-glycine gels and fractionated at 150 V for 1.5 h, followed by transfer to a PVDF membrane for 2 h at 300 mA and incubation with the antibodies listed above. For quantitation of western blots, chemiluminescence images were analysed in the ImageLab software where intensity of bands was noted from at least 3 blots taken from 3 experiments.

### GST pulldown assay

Cells were lysed (20 mM Tris-HCl pH 7.5, 150 mM NaCl, 1% Triton X-100) and precleared by incubating with 30 μL GST beads for 1 h to reduce non-specific binding. We then added 1 mg of the precleared lysate to 5 μg of GST-tagged protein and 20 μL glutathione Sepharose resin, and incubated at 4 °C for 2 h on a rotary shaker. The resin is then washed (20 mM Tris-HCl pH 7.5, 300 mM NaCl, 1% Triton X-100), boiled with SDS sample buffer and analyzed by western blot as described above.

### PR-Ub assay

5mM MBP/His tagged p62 were incubated at 37 °C for 1 h with 25 mM of purified untagged ubiquitin, 1 mM NAD^+^ and 1–2 mM SdeA in 50 mM Tris-HCl (pH 7.5) and 50 mM NaCl. The reaction mixture was processed as described for western blotting above, and probed with antibodies specific for ubiquitin and GST. PR-Ub-specific deubiquitination assays were performed by incubating PR-Ub proteins with 1 μg of GST-DupA at 37 °C for 1 h in a buffer containing 50 mM Tris-HCl (pH 7.5) and 150 mM NaCl.

### Identification of PR-Ub sites

PR-Ub sites were identified by ETD-MS as previously described (Leidecker et al., 2016, Liu et al., 2021). MBP-tagged p62 was modified by SdeA *in vitro* then denatured in 0.1 M Tris-HCl (pH 7.5) containing 8 M urea. The samples were washed three times in 200 µL of the same buffer in a 30-kDa Amicon Ultra 0.5-mL centrifugal filter (Merck) to remove free ubiquitin. The eluted protein was washed twice in 50 mM ammonium bicarbonate (pH 7.5) before digestion using a 1:50 ratio of trypsin the same buffer for 6 h. The peptides were desalted on a C18 column, followed by LC-MS/MS analysis. The spectra were collected and deconvoluted, and high-resolution ETD-MS/MS spectra were manually inspected to verify the sequence.

### Legionella infection

*Legionella pneumophila* strains were obtained from Dr. Zhao-Qing Luo (Purdue University) and were grown for 3 days on *N*-(2-acetamido)-2-amino-ethanesulfonic acid (ACES)-buffered charcoal (BCYE) extract agar, at 37 °C, followed by growth for 20 h in CYE medium. Bacterial cultures (OD_600_ = 3.2–3.6) were used to infect HEK 293T, HeLa cells with a multiplicity of infection (MOI) of 10, and A549 cells with a MOI of 2. For 12-h infection experiments, we used a MOI of 1. HEK 293T and HeLa cells were transfected with CD32 16 h before infection to facilitate bacterial entry.

### Intracellular replication of *Legionella*

A549 cells growing in 35-mm dishes were infected with Legionella strains at a MOI of 1. The infection was allowed to proceed for 90 min in antibiotic-free medium before switching to medium containing gentamycin for 60 min to kill the remaining extracellular bacteria. The cells were then lysed in 0.4% saponin immediately or after 24 or 48 h to release intracellular bacteria. Lysates were spotted onto BCYE plates at 1:10 and 1:100 dilutions. The intracellular bacterial load was assessed by counting bacterial colonies formed after 48 h. The number of colony forming units (cfu) was calculated using the formula cfu/mL = (number of colonies × dilution factor) / volume of sample plated (mL). The fold-increase in cfu was calculated by normalizing the cfu at 24 or 48 h to that determined immediately after lysis.

### Confocal microscopy and image analysis

Confocal images were observed using a Zeiss LSM780 microscope system fitted with a 63× 1.4 NA oil-immersion objective as well as argon and helium–neon ion lasers for the excitation of GFP (488 nm) and RFP (546 nm), respectively. Images were analyzed in FIJI to determine recruitment of proteins to bacteria. For statistical analysis, atleast 50 cells were analysed from 3 independent experiments.

For measurement of intracellular ROS, 5µM Cell ROX green reagent (C10444, Thermofisher Scientific) was loaded to bacteria infected A549 cells for 30min followed by fixation using 4% paraformaldehyde. Cells were immunostained using anti-Legionella antibody and imaged using the 488nm laser line and 40X objective of the Zeiss LSM780 system. All imaging parameters (laser power, magnification and gain settings) were kept identical to compare intensities between samples. Images were analysed in FIJI by measuring 488nm (green) signal collected from the infected cells marked manually as multiple region of interest (ROIs). 100 cells per set were analysed per set taken from 3 experiments.

### Redox GFP measurements

Redox GFP(roGFP) targeted to the mitochondrial matrix or cytosolic redox GFP was transfected in A549 cells 24h prior to bacterial infection. Cells were infected with Legionella for different lengths of time and imaged using confocal imaging. For microscopy, excitation wavelengths of 365nm, and 488nm were used sequentially. In both cases emission signals were collected uusing the GFP filter (emission maxima 510nm). Oxidised roGFP is excited at 365nm while the reduced form of is excited at 470nm. Z-stacks with 0.5µm slices were imaged for at least 60 cells taken from 3 experiments and analysed in FIJI as before (Vevea et al., 2013). Briefly, images were converted to 32 bit, background subtracted, thresholded and the ‘image calculator was used to divide the signal from oxidised roGFP (365nm excitation) by reduced roGFP (Excitation 488nm). This ratio was calculated from 60 cells and plotted in the online tool for boxplot generation boxplotR (http://shiny.chemgrid.org/boxplotr/)

### Luciferase activity assay

To analyze the induction of Nrf2 induced genes, a luciferase reporter assay was used in A549 cells. In brief, an expression construct containing the luciferase ORF and the NRF2 promoter with antioxidant response element (Nrf2-ARE) was transfected. In total, 100 ng plasmid was used per one well of a 12-well dish. All transfections were performed in triplicate and the average of three experiments is shown in figures. Sixteen hours after transfection, cells were infected with bacteria for indicated timespans. Luciferase expression was measured using the Luciferase Reporter Assay System (Promega). Fold change was calculated by taking untreated cells as 1.

### Nuclear fractionation

A549 cells from a confluent 60 mm dish transiently transfected Myc-Nrf2. 16 h later they are infected with Legionella strains for 6 h. Cells were lysed in 300 µL of hypotonic buffer [10 mM HEPES (pH 7.4), 2 mM MgCl2, 25 mM KCl, 1 mM DTT, 1 mM PMSF, and protease inhibitor cocktail], kept on ice for 30 min followed by syringe lysis, then 125 µL of a 2M sucrose solution was added dropwise, followed by centrifugation at 1000g for 15 min. The supernatant was saved as the cytosolic fraction. The pellet was washed twice in wash buffer [10mM HEPES (pH 7.4), 2mM MgCl2, 25mM KCl, 250mM sucrose, 1mM DTT, 1 mM PMSF, and protease inhibitor cocktail] and saved as the nuclear fraction.

### Nrf2 ubiquitination assay

A549 cells expressing Myc-Nrf2 were infected with Legionella for 6h and treated with 20µM MG132 2h prior to lysis. Cells were lysed in 50mM Tris (pH 7.5), 150mM NaCl, 2% SDS and 10 mM *N*-ethylmaleimide. Lysates were were diluted fivefold in buffer containing 50 mM Tris-HCl, pH 7.5, 150 mM NaCl and 1% Triton X-100. Lysates were incubated with myc agarose resin for two hours at 4°C, followed by washing in wash buffer containing 50 mM Tris-HCl, pH 7.5, 400 mM NaCl and 1% Triton X-100. Samples were analyzed by SDS gel electrophoresis, transferred to PVDF membranes, and immunoblotted with the anti-Ub, and Myc antibodies.

### Quantification of cellular RNA

For real time PCR, total RNA was isolated using Trizol according to the manufacturer’s instructions. In total, 1 μg RNA was converted to cDNA using random hexamer primer using Thermo cDNA synthesis kit. RT-PCR was performed using EvaGreen (Biotium, Catalog no: 89138-982) and the Biorad CFX Connect Real-time PCR system. Gene expression was compared using C_t_ values and the results were calculated using ^ΔΔ^C_t_ method with normalization to the average expression level of 18S rRNA. Forward (F) and reverse (R) primers used are as follows:

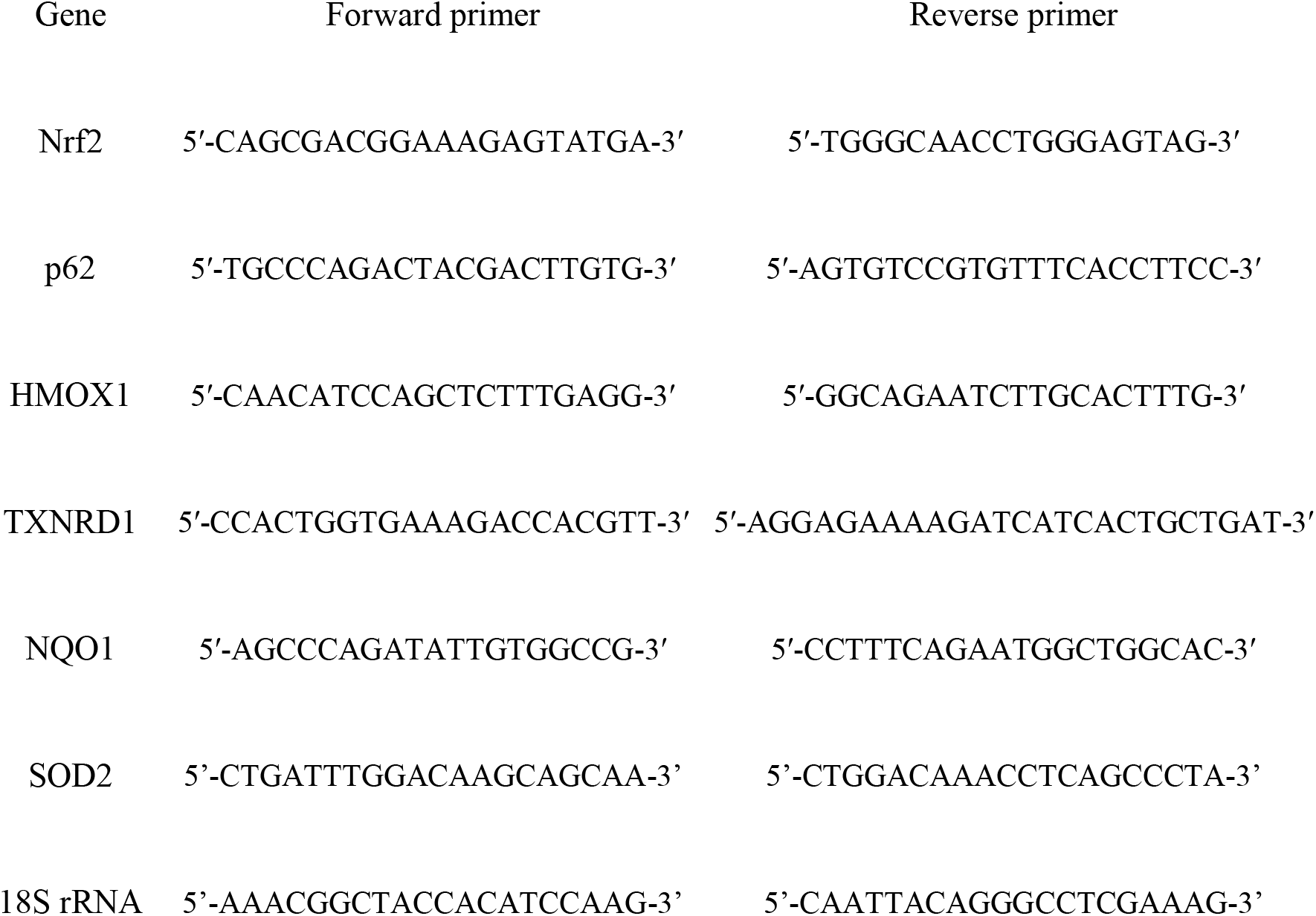

### RNA sequencing

50µg RNA was isolated using the Qiagen mRNA isolation kit. Quality of RNA was assessed by checking on an agarose gel to see the intactness of the 28S, and 18S rRNA bands and absorbance at 260nm and 280nm was measured by Nanodrop. Absorbance(260nm:280nm) was >1.8, and Absorbance(260nm:230nm)>2.0. Isolated RNA was used for next generation RNA sequencing using Illumina platform. Data was analyzed by ROSALIND® (https://rosalind.bio/), with a HyperScale architecture developed by ROSALIND, Inc. (San Diego, CA). Reads were trimmed using cutadapt (Martin et al, 2011). Quality scores were assessed using FastQC (Andrews, 2010). Reads were aligned to the Homo sapiens genome build GRCh38 using STAR (Dobin et al, 2013). Individual sample reads were quantified using HTseq (Anders et al., 2015) and normalized via Relative Log Expression (RLE) using DESeq2 R library (Love et al., 2014). Normalized expression of genes between 4 replicates each of WT, ΔS and non-infected cells were compared. Students t test (2 tailed type 3) was used to calculate p values. Volcano plots were generated to visualise differential gene expression between WT and ΔS Legionella infection.

### Sample preparation for mass spectrometry

A549 cells were infected with Legionella for 6h in presence of tBHQ (10µM) or ML385 (10µM). Cells were lysed in SDS-lysis buffer (50 mM Tris, 2% SDS, 10 mM TCEP, 40 mM chloroacetamide, protease inhibitor cocktail, phosphatase inhibitor) heated to 95 °C for 10 min. Proteins were precipitated using methanol-chloroform precipitation and resuspended in 8 M urea. Isolated proteins were digested with 1:50 w/w LysC (Wako Chemicals, cleaves at the carboxylic side of lysine residue) and 1:100 w/w trypsin (Promega, Sequencing-grade; cleaves at the carboxylic side of lysine and arginine residues) overnight at 37 °C after dilution to a final urea concentration of 1 M. Digests were then acidified (pH 2–3) using trifluoroacetic acid (TFA). Peptides were purified using C18 SepPak columns (Waters). Desalted peptides were dried and 25 μg of peptides was resuspended in TMT-labeling buffer (200 mM EPPS pH 8.2, 20% acetonitrile). 160µg Peptides were subjected to TMT labeling with 1:2 peptide TMT ratio (w/w) for1 h at room temperature. The labeling reaction was quenched by addition of hydroxylamine to a final concentration of 0.5% and incubation at room temperature for an additional 15 min.

### Mass spectrometric data acquisition

Samples were pooled in equimolar ratio and subjected to High pH Reversed-Phase Peptide Fractionation kit (Thermo Fisher Scientific) following the manufacturer’s instructions. Eluted fractions were dried in SpeedVac and peptides were resuspended in 3% acetonitrile/0.1% TFA for liquid chromatography–mass spectrometry. All mass spectrometry data was acquired in centroid mode on an Orbitrap Fusion Lumos mass spectrometer hyphenated to an easy-nLC 1200 nano HPLC system with a nanoFlex ion source (Thermo Fisher Scientific). Resuspended peptides were separated on an Easy nLC 1200 (Thermo Fisher Scientific) and a 22-cm-long, 75-μm-innerdiameter fused-silica column, which had been packed in house with 1.9-μm C18 particles (ReproSil-Pur, Dr Maisch) and kept at 45 °C using an integrated column oven (Sonation). Peptides were eluted by a nonlinear gradient from 8 to 60% acetonitrile over 155 min followed by an increase to 95%B in 1 min, which was held for another 10 min. Full-scan MS spectra (350–1400 m/z) were acquired at a resolution of 120,000 at m/z 200, a maximum injection time of 100 ms, and an AGC target value of 4 × 10^5^. Up to 20 most intense precursors, with charge state in between 2 and 5, were isolated using a 0.7 Th window. MS/MS spectra were acquired with a maximum injection time of 50 ms, AGC target value of 1.5 × 10^4^, and fragmented using CID with a normalized collision energy of 35%. SPS-MS3 scans were done on the ten most intense MS2 fragment ions having an isolation window of 0.7 Th (MS) and 2 m/z (MS2). Ions were fragmented using NCE of 50% and analyzed in the orbitrap with the resolution of 50,000 at m/z 200, scan range 110–500 m/z, AGC target value 1.5 × 10^5^, and a maximum injection time of 120 ms.

### Mass spectrometric data analysis

Raw mass spectrometry data were analyzed with Proteome Discoverer (v.2.4, Thermo Fisher Scientific) using Sequest HT as a search engine and performing recalibration of precursor masses by the Spectrum RC-node. Number of missed cleavages permitted was 2. Mass tolerance of precursor ions is 7 ppm and mass tolerance of fragment ions is 0.5 Da. Fragment spectra were searched against the human reference proteome (“one sequence per gene,” 20,531 sequences, version March 2020) and protein sequences SARS-CoV-2 (14 sequences, version February 2020) downloaded from Uniprot in March 2020, as well as common contaminants as included in “contaminants.fasta” provided with the MaxQuant software (version 1.6.11). Static modifications were TMT at the peptide N-terminus and lysines as well as carbamidomethyl at cysteine residues, dynamic modifications were set as oxidation of methionine and acetylation at the protein N-terminus. Matched spectra were filtered with Percolator, applying a false discovery rate of 1% on peptide spectrum match and protein level. Reporter intensities were normalized to the total protein intensities in Proteome Discoverer, assuming equal sample loading and additionally by median normalization using the NormalyzerDE package. Statistically significant changes between samples were determined in Perseus (v.1.6.6.0). Data set was first filtered for contaminants and biological replicates were grouped as one. Proteins were further filtered for two valid values present in at least one group. Missing values were imputated from normal distribution of data using default settings. Significant candidates were chosen using two-sided *t* test with error-corrected *p*-values (0.01. FDR) and log2(fold change) value minimum of ±0.5. To avoid false-positive protein identification, ≥2 unique peptides identified within a single protein were used for further analysis. Network and gene ontology analysis was performed with statistically significant hits using the online Metascape software

## Conflict of interest

The authors declare that they have no conflicts of interest with the contents of this article.

## Acknowledgments

We would like to thank Florian Bonn for his help with the DSF mass spectrometric determination of PR-Ub sites.

## Author contributions

R. M. and I. D. conceptualization; R.M., S.K.K, A. G, A.B. data curation; I. D. funding acquisition; R. M., S.K.K., A.G, A.B, D. S., D.S, and I. D. investigation; R. M., A. B., R. R, D.S, T.C, and M.M. methodology; I. D. project administration;. I.M and I. D. resources; I.M and I. D. supervision; R. M. and I. D. validation; R. M. visualization; R. M. writing-original draft; R. M., S.K.K, A.G, A.B, R.R, D.S, M.M, T.C, I.M and I.D writing-review and editing.

## Funding and additional information

I.D. acknowledges funding from the Deutsche Forschungsgemeinschaft (DFG, German Research Foundation) Project-ID 323732846-LYSFOR2625 and Project-ID 25913077-SFB1177. D. S. was supported by the National Research Foundation of Korea (NRF) grant funded by the Korea government (MSIT) (No. 2021R1C1C1003961) and the Yonsei University Research Fund of 2021 (2021-22-0050). RM acknowledges Humboldt Stiftung for postdoctoral fellowship, SKK is funded through EMBO postdoctoral fellowship (ALTF 199-2021).

## Supplementary Figure Legends

**Figure S1:**
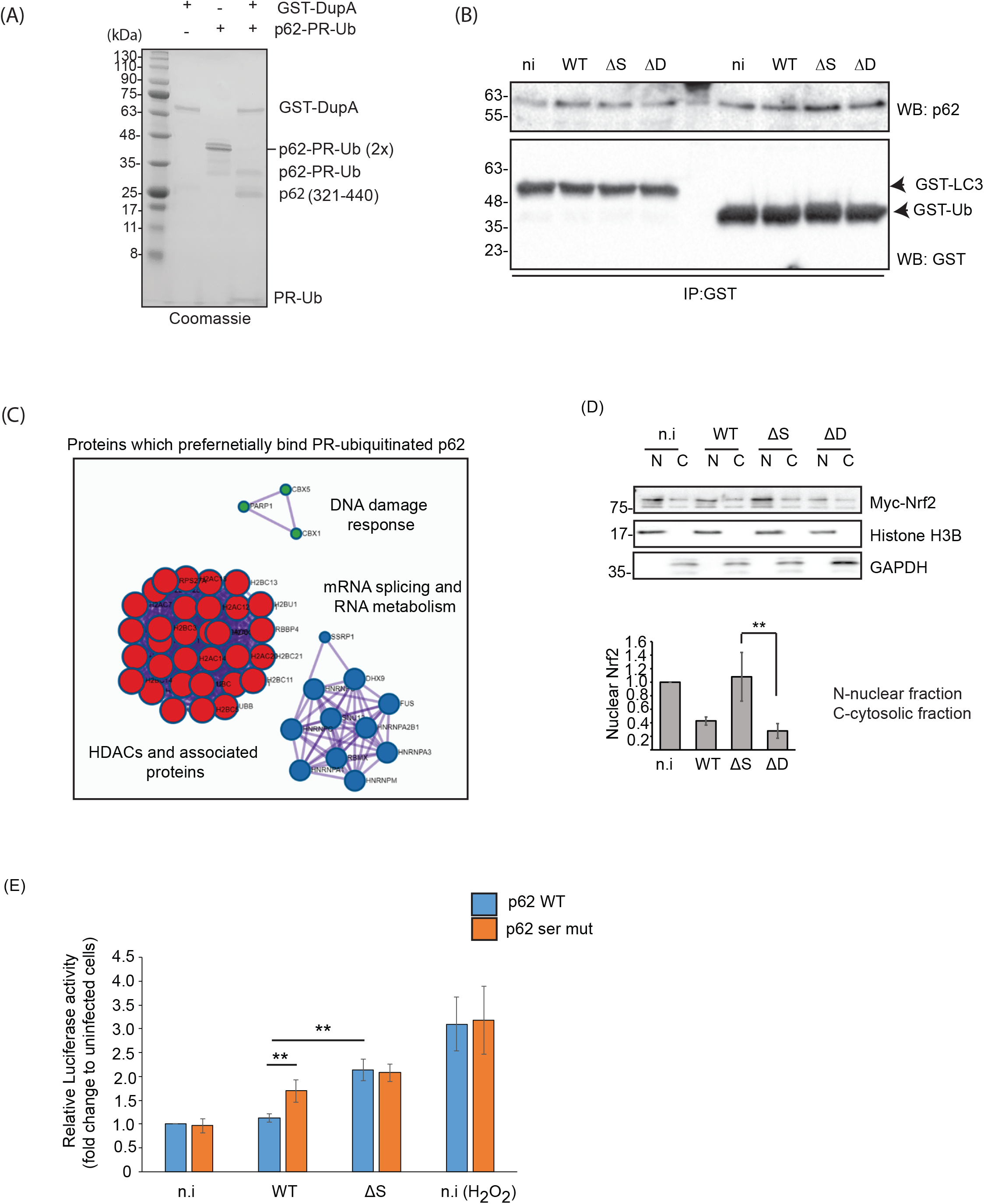
PR-Ub of p62 *in vitro* does not alter its interaction with LC3 and Ub but regulates Nrf2 dependent gene expression. A. P62 (amino acids 321-400) is PR-ubiquitinated *in vitro* by incubating it with SdeA, Ub and NAD. Treatment with GST-DupA removes the PR-Ub from p62 B. GST tagged LC3, Ub were incubated with lysate from HEK293T cells infected with mentioned Legionella strains followed by GST pulldown and immunoblotting with GST, and p62 antibodies. C. GST-p62(amino acids 321-400) of p62 purified from E. coli, was PR-ubiquitinated it in an *in vitro* reaction or left unmodified and incubated it with cell lysate from HEK293T cells infected with WT Legionella in a GST pull down assay, followed by mass spectrometric identification of interactors. List of proteins significantly enriched in the PR-Ub modified p62 pulldown were put in the online Metscape software to identify enriched pathways. D. A549 cells expressing Myc-Nrf2 were infected with Legionella strains for 6 h and fractionated into nuclear and cytosolic fractions followed by immunoblotting of fractions with Nrf2. HistoneH3B and GAPDH were used as nuclear and cytosolic loading controls respectively. Graph represent 3 experiments; error bars indicate standard deviation. ** 0.01>p≥0.001 E. Relative luciferase-ARE activity in A549 cells expressing wt or PR-Ub deficient p62 and infected with Legionella strains for 6 h (n.i-not infected, WT-wild-type Legionella, ΔS-ΔSidE Legionella, ΔD-ΔDupA/B Legionella)

**Figure S2:**
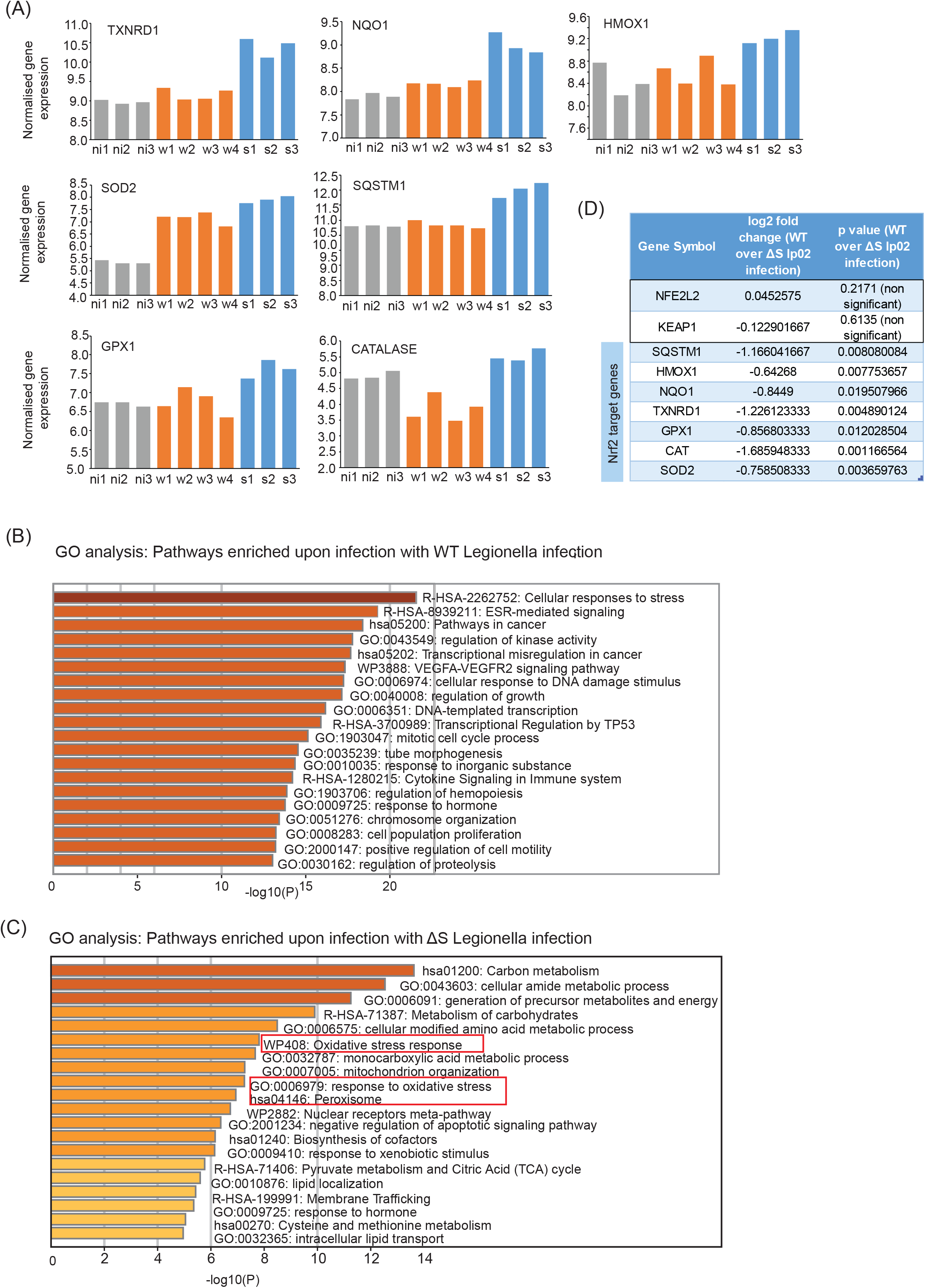
Expression of Nrf2 targets from RNA sequencing. A. Normalized gene expression of Nrf2 target genes in replicates of non-infected (ni1-ni3), WT (w1-w4) or ΔS Legionella(s1-s3) infected cells as measured by next gen sequencing B. GO analysis of genes whose expression are significantly upregulated in WT Legionella infection C. GO analysis of genes whose expression are significantly upregulated in ΔS Legionella infection D. Nrf2 target genes identified by RNAseq and enriched in WT over ΔS Legionella infection.

**Figure S3:**
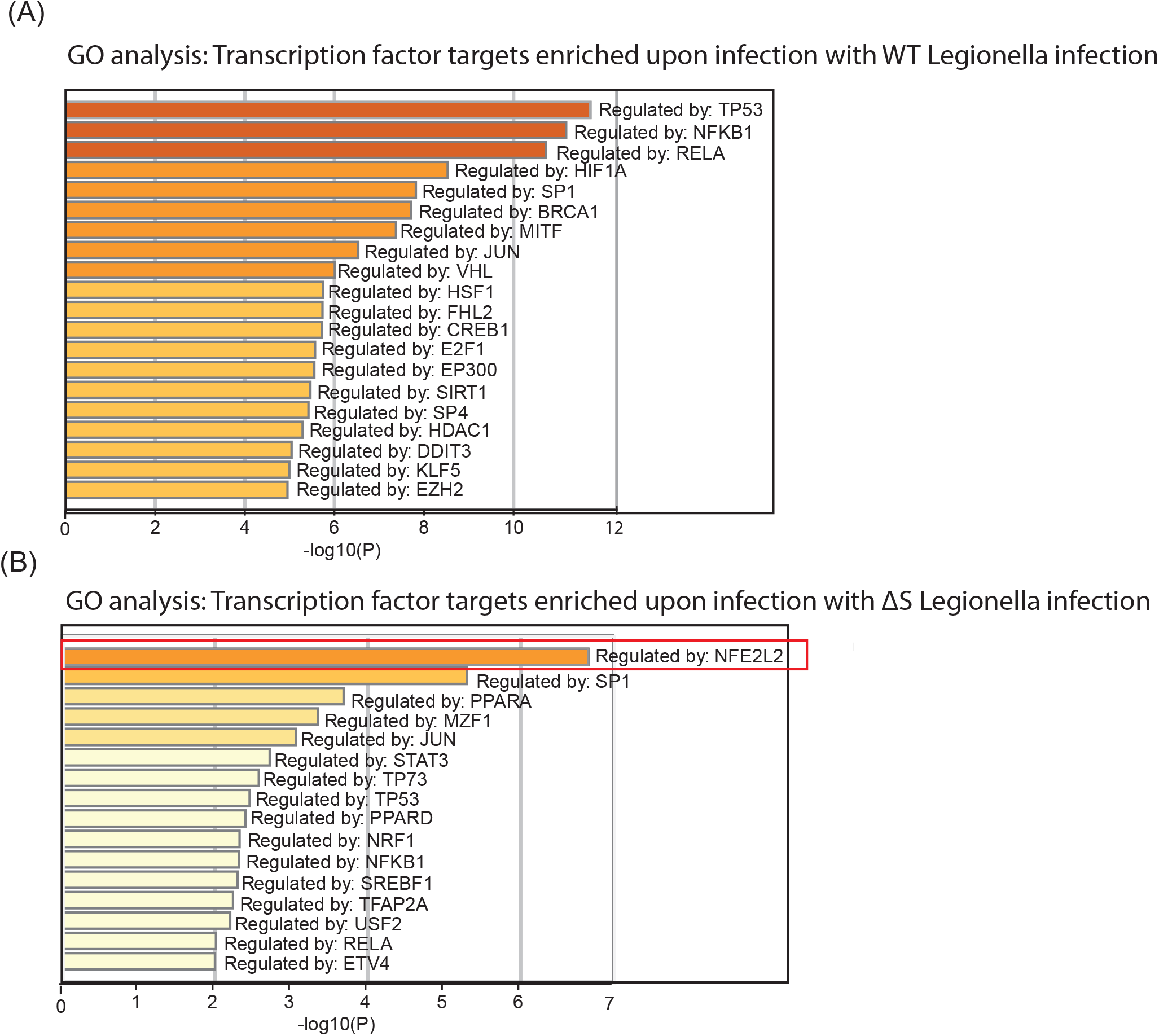
Transcriptional pathways differentially regulated by PR-Ub from RNAseq. A. Transcriptional pathways significantly upregulated in WT Legionella infection. B. Transcriptional pathways significantly upregulated in ΔS Legionella infection.

**Figure S4:**
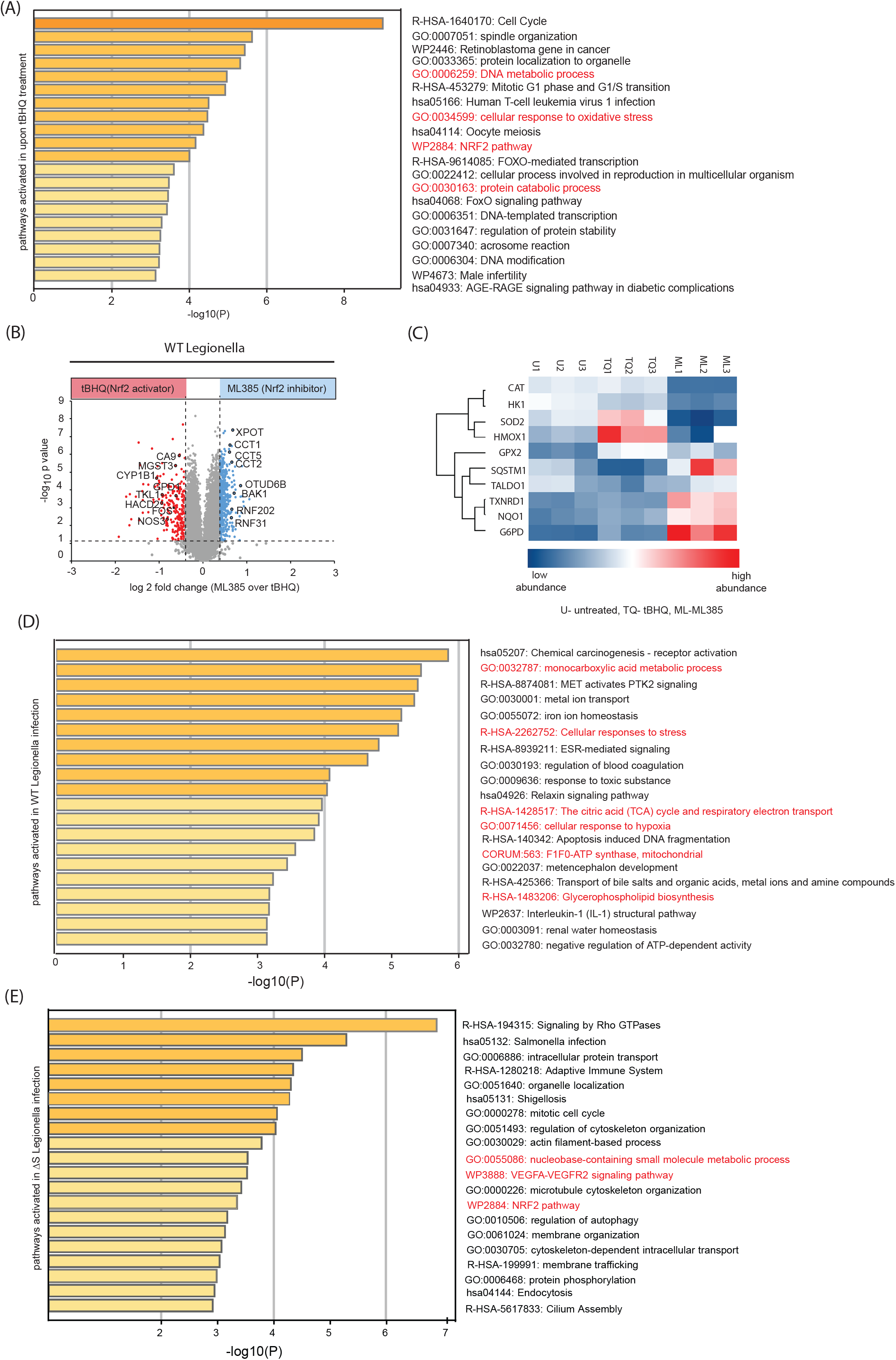
Nrf2 dependent proteome changes in Legionella infection. A. A549 cells were treated with 10µM tBHQ or ML385 to activate and inhibit Nrf2 dependent gene expression respectively. Cells were lysed after 8 h of treatment followed by protein extraction, peptide enrichment and analysis of the proteome by mass spectrometric analysis. Proteins enriched upon tBHQ treatment were subjected to GO analysis. Pathways related to metabolism and oxidative stress are marked in red. B. Volcano plot showing changes in the proteome of A549 cells infected with WT Legionella in presence of 10µM tBHQ or 10µM ML385 for 8 h. C. Heatmap showing levels of Nrf2 pathway proteins in the experiment indicated in panel b. D. GO analysis of proteins enriched at 8 h.p.i of WT Legionella infection in presence of 10µM tBHQ. Pathways related to metabolism and oxidative stress are marked in red. E. GO analysis of proteins enriched at 8 h.p.i of ΔS Legionella infection in presence of 10µM tBHQ. Pathways related to metabolism and oxidative stress are marked in red.

**Figure S5:**
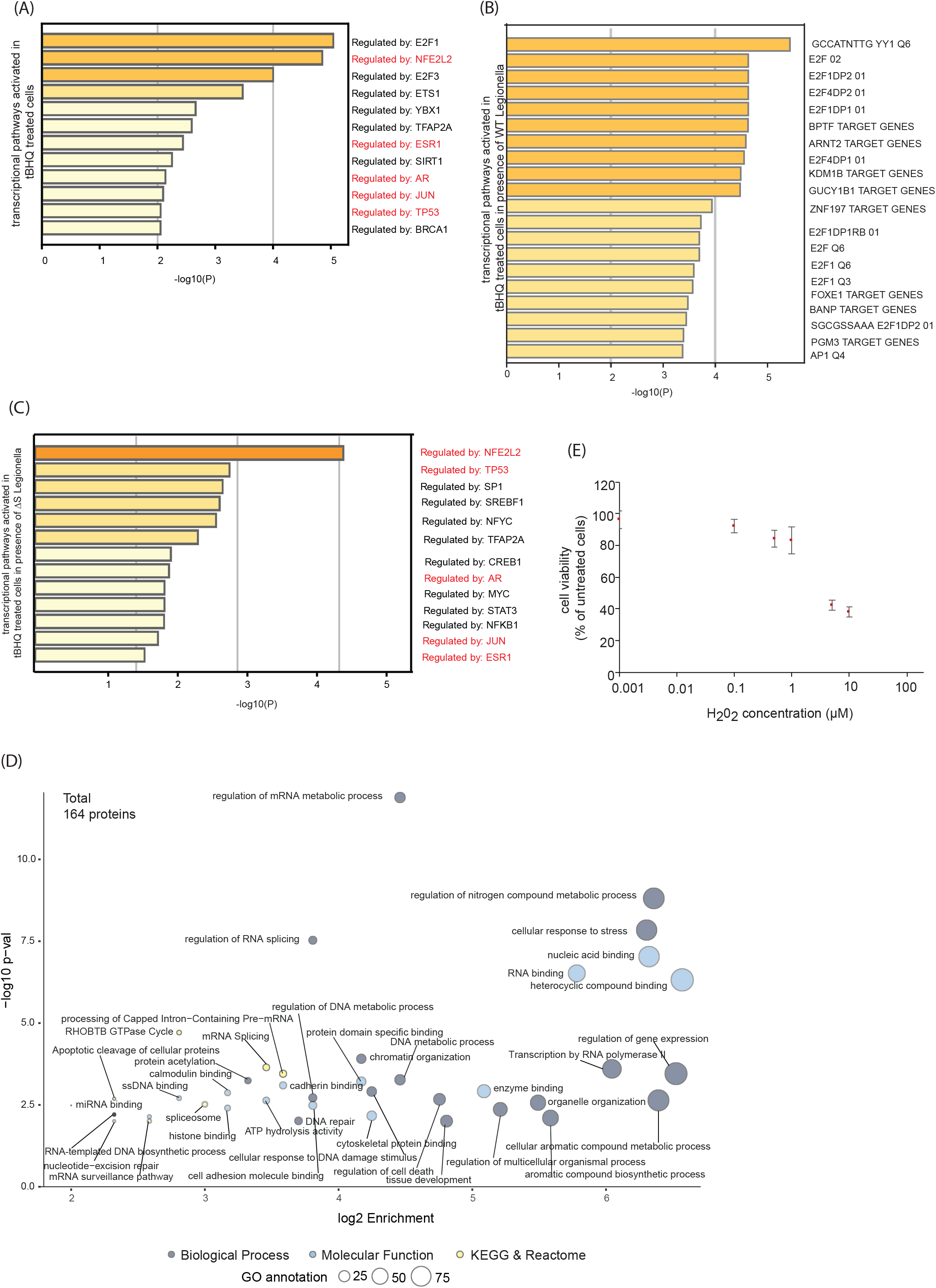
Transcriptional networks and phosphosites altered in the proteome in Legionella infection. A. TRRUST analysis of proteins enriched upon treatment with 10µM tBHQ for 8 h. B. TRRUST analysis of proteins enriched at 8 h.p.i of WT Legionella infection in presence of 10µM tBHQ C. TRRUST analysis of proteins enriched at 8 h.p.i of ΔS Legionella infection in presence of 10µM tBHQ D. GO analysis of the 164 proteins differentially regulated by phosphorylation during ΔS Legionella infection. E. A549 cells were treated with different concentrations of H_2_O_2_ for 48 h and cell viability was assessed by a MTT assay. (transcriptional networks regulated by tBHQ which are shared between uninfected and ΔS Legionella infection are marked in red)

## Notes

### Competing Interest Statement

The authors have declared no competing interest.

